# uniLIVER: a Human Liver Cell Atlas for Data-Driven Cellular State Mapping

**DOI:** 10.1101/2023.12.09.570903

**Authors:** Yanhong Wu, Yuhan Fan, Yuxin Miao, Yuman Li, Guifang Du, Zeyu Chen, Jinmei Diao, Yu-Ann Chen, Mingli Ye, Renke You, Amin Chen, Yixin Chen, Wenrui Li, Wenbo Guo, Jiahong Dong, Xuegong Zhang, Yunfang Wang, Jin Gu

## Abstract

The liver performs several vital functions such as metabolism, toxin removal and glucose storage through the coordination of various cell types. The cell type compositions and cellular states undergo significant changes in abnormal conditions such as fatty liver, cirrhosis and liver cancer. As the recent breakthrough of the single-cell/single-nucleus RNA-seq (sc/snRNA-seq) techniques, there is a great opportunity to establish a reference cell map of liver at single cell resolution with transcriptome-wise features. In this study, we build a unified liver cell atlas uniLIVER by integrative analyzing a large-scale sc/snRNA-seq data collection of normal human liver with 331,125 cells and 79 samples from 6 datasets. Besides the hierarchical cell type annotations, uniLIVER also proposed a novel data-driven strategy to map any query dataset to the normal reference map by developing a machine learning based framework named LiverCT. Applying LiverCT on the datasets from multiple abnormal conditions (1,867,641 cells and 439 samples from 12 datasets), the alterations of cell type compositions and cellular states were systematically investigated in liver cancer.

## Main

The liver is a major metabolic organ, which performs many essential physiological functions, including toxin removing, albumin and bile production, glucose and amino acid processing, and vitamin storage, etc. For humans, hepatocytes occupy about 80% liver volume and cholangiocytes, immune cells and stromal cells consist of the majority of the remaining part. These cells are organized into hexagonal hepatic lobules as the basic functional units of liver. As the recent breakthrough of the single-cell/single-nucleus RNA-seq (sc/snRNA-seq) techniques^1,2^, there is a great opportunity to establish a reference cell map of liver at single cell resolution with transcriptome-wise features. Besides, the reference map is very useful for studying the altered cell type compositions and cellular states under diverse physiological and pathological conditions in liver, such as acute injury, virus infection, cirrhosis and cancer.

In this study, we collected 79 normal human liver samples from 6 datasets^3–7^, and 439 abnormal or disease samples from 12 datasets^4,8–18^. Based on this collection, we hierarchically annotated 63 cell types/subtypes and the hepatocytes in 4 different lobular regions/zones of normal liver, and then constructed an integrated and data-driven human liver cell atlas uniLIVER. Beyond the traditional cell atlases mainly providing comprehensive cell type annotations, uniLIVER also aims at establishing a novel data-driven strategy to map any query dataset to the normal reference map. Analogy to the genome sequence mapping, we proposed a concept for cell type or cellular state mapping: the query cells are computationally “mapped” to the reference map based on gene expression features. Those cells whose gene expressions are dissimilar to any annotated cell subtype in the reference are defined as “variant” state cells (analogy to nucleotide variants in genome sequence analysis). The “variant” states are broadly categorized into two types, the “deviated” state and the “intermediate” state: the “deviated” state means that the gene expressions of the query cells are shifted from a single cell type, and the “intermediate” state means that the gene expressions of the query cells located between any two reference cell types. We developed LiverCT, a machine learning based liver **<u>C</u>**ell-**<u>T</u>**ype mapping method using the annotated reference data, to achieve the task of cellular state mapping in liver.

We applied LiverCT on the collected abnormal liver datasets. Results show that almost all types of cells are strongly “deviated” from their normal states in hepatocellular carcinoma (HCC) and the deviated scores of T cells are positively correlated with the stress response pathway signatures. Interestingly, the results also show that the hepatic stellate cells and granulocytes (mainly neutrophils) are highly deviated in adjacent non-tumor tissues. For the “intermediate” state analysis, it was found that the cancer cells with high intermediate scores have strongly up-regulated glycolysis and hypoxia pathways. Also, the up-regulated genes of those cells are significantly overlapped with poor prognosis genes. Another interesting task is to analyze the zonation tendency of the HCC tumor cells by mapping them to the hepatocyte states in different lobular zones. We found that the zonation tendency is highly associated with the expression of multiple malignant signatures and the composition of multiple immune and stromal cell types.

uniLIVER tends to establish a new framework of cell atlas by introducing machine learning into the traditional data portal only design. Both the reference cell map (as three portraits similar to hECA^19^) and the computationally cellular state mapping tool LiverCT are freely available via a web-based database.

## Results

### Overview of the human liver cell atlas uniLIVER

We collected scRNA-seq data from 6 datasets, which include 79 samples from 42 donors (Table 1). After stringent quality control, 331,125 cells were integrated and annotated hierarchically to form the normal reference map. Population and clinical information, such as age and gender, were also collected if available (Extended Data Fig. 1a, Supplementary Table 1). Besides, we also curated 1,867,641 cells from diverse abnormal or disease samples including cirrhosis, hepatocellular carcinoma (HCC), intrahepatic cholangiocarcinoma (ICC), combined hepatocellular and intrahepatic cholangiocarcinoma (CHC) and some liver metastases (Table 1).

**Table 1.** The collected datasets in uniLIVER.

| Study | No. of<br>donors | Gender<br>(M:F) | Age<br>(yr) | Sequencing<br>method | Final<br>#cells | Source |
| --- | --- | --- | --- | --- | --- | --- |
| Aizarani et al. 2019<sup>3</sup> | 9 | n/a | n/a | mCel-Seq2 | 9,466 | GSE124395 |
| Ramachandran et al. 2019/healthy<sup>4</sup> | 5 | 4M:1F | n/a | 10X | 34,601 | GSE136103 |
| Guilliams et al. 2022<sup>5</sup> | 34 | 19M:15F | 28-77 | 10X | 167,510 | GSE192742 |
| Payen et al. 2021<sup>6</sup> | 2 | 1M:1F | 3-39 | 10X | 26,685 | GSE158723 |
| Andrews et al. 2021<sup>7</sup> | 5 | 2M:3F | 18-60 | 10X | 59,977 | GSE185477 |
| Gu et al. 2022 | 4 | 2M:2F | 47-66 | 10X | 32,886 | In house |
| Losic et al. 2020<sup>8</sup> | 2 | 1M:1F | 66-67 | 10X | 49,674 | GSE112271 |
| Lu et al. 2022<sup>9</sup> | 10 | 9M:1F | 48-65 | 10X | 71,915 | GSE149614 |
| Ma et al. 2021<sup>10</sup> | 37 | 23M:13F | 35-81 | 10X | 48,318 | GSE116113 |
| Massalha et al. 2020<sup>11</sup> | 6 | 2M:4F | 40-74 | MARS-seq | 4,691 | GSE146409 |
| Sun et al. 2021<sup>12</sup> | 18 | 17M:1F | 42-76 | MIRALCS | 16,498 | CNP0000650 |
| Zhang et al. 2019<sup>13</sup> | 10 | 9M:1F | 32-84 | 10X/SMART<br>-seq2 | 73,261 | GSE140228 |
| Zhang et al. 2020<sup>14</sup> | 6 | 4M;2F | n/a | 10X | 37,814 | GSE138709/<br>GSE142784 |
| Zheng et al. 2017<sup>15</sup> | 6 | 4M;2F | 26-64 | SMART-seq2 | 5,063 | GSE98638 |
| Ramachandran et al. 2019/cirrhosis<sup>4</sup> | 5 | 3M:2F | n/a | 10X | 26,279 | GSE136103 |
| Xue et al. 2022<sup>16</sup> | 124 | 94M:30F | 31-88 | 10X | 1,337,829 | PRJCA007744 |
| Ma et al. 2022<sup>17</sup> | 7 | n/a | n/a | 10X | 112,506 | GSE189903 |
| Liu et al. 2023<sup>18</sup> | 6 | 6F | 48-64 | 10X | 83,793 | skrx2fz79n |

Then, the atlas tends to “map” the cells from disease samples to the normal reference map based on gene expression similarities. To find the “variant” state cells in disease samples, we developed LiverCT, a machine learning based **<u>C</u>**ell-**<u>T</u>**ype mapping method that can distinguish “deviated” states and “intermediate” states and can also classify the abnormal hepatocytes in HCC into P-like (P: Portal/Periportal) state, M-like (M: Mid) state and C-like (C: Central) state (Fig. 1). One unique aspect of LiverCT is that it adopts the concept of genomic variant analysis to identify and analyze the “variant” states of cells based on their expression patterns. Firstly, LiverCT embedded the cells in the normal reference map into a latent space. In the latent space, we trained a hierarchical classifier using ensemble learning to predict the cells’ type labels. This was followed by another one-class classifier to identify the margin of a normal cell type and a one-vs-one classifier to identify the boundary between any two cell types. Further, if a cell was predicted as hepatocyte, we utilized an additional classifier to distinguish its lobular zonation tendency. When applied to query datasets, LiverCT embedded the query cells into the pre-calculated latent space of the normal reference by scArches^20^ to remove the batch effects. After the cell type prediction, a “deviated” score was calculated to measure the degree of deviation from the corresponding normal cell type and an “intermediate” score was calculated to measure the degree to which the cell was in the middle of top two predicted cell types. (Methods, Extended Data Fig. 1b).

**Fig. 1.**
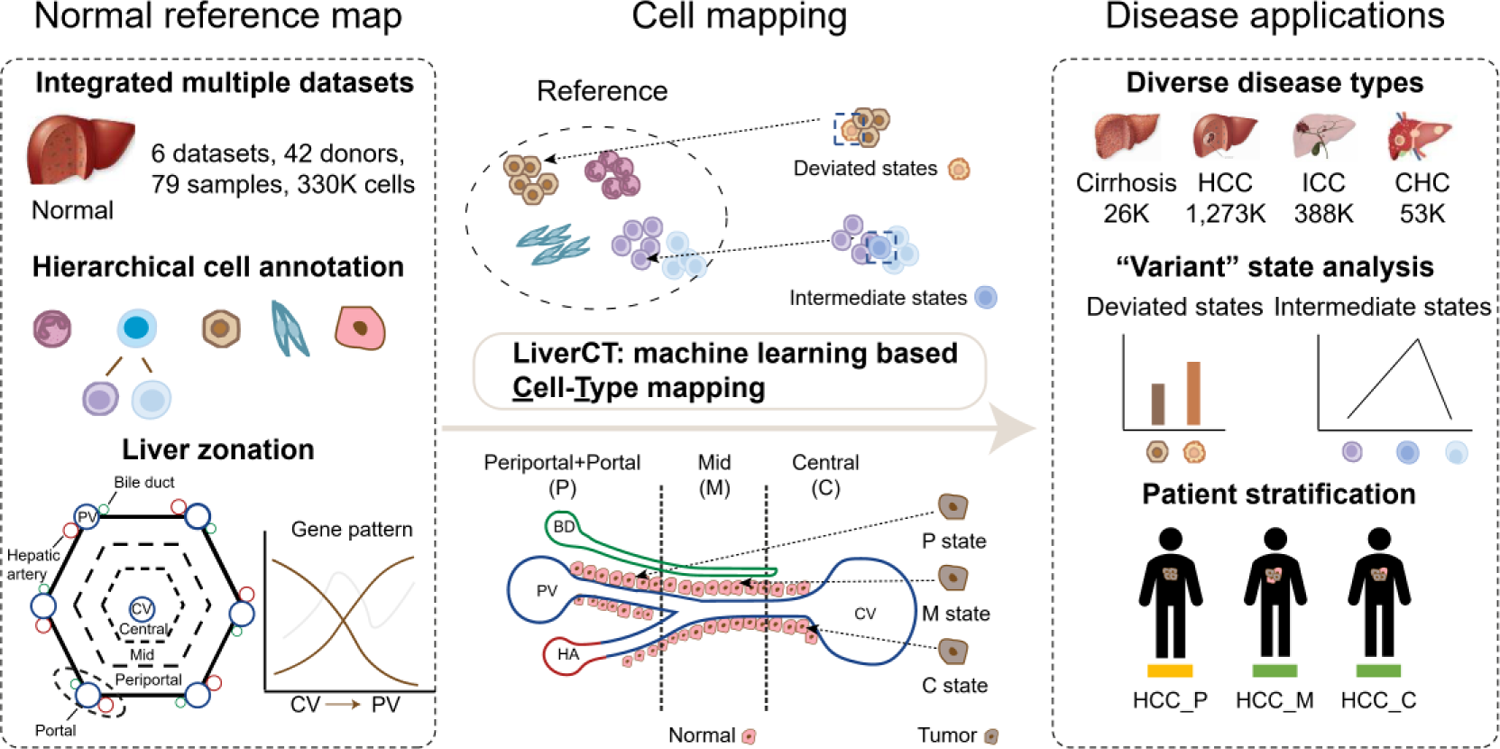
uniLIVER overview. The normal reference by hierarchically annotating the cell types and zonation tendency via an integrative analysis of multiple sc/snRNA-seq datasets (left). The machine learning based Cell-Type mapping method named LiverCT can identified the cells with deviated or intermediate states and calculate the hepatocyte zonation tendency of any query dataset (middle). LiverCT was applied for liver cancer and other abnormal conditions (rights).

### Hierarchical annotations of the normal liver cells

A unified normal reference map needs in-depth and harmonized annotations. Currently, there exists some inconsistency in cell type definition across different studies. To harmonize the cell type labels from different datasets, firstly we built a unified hierarchical annotation framework (uHAF^19^) for liver (Fig. 2a). The major cell type annotations provided by the original references were harmonized into 8 major cell types (Level 1) in uniLIVER (Supplementary Table 2). In cases where the annotations were not provided, the datasets were annotated manually. Using the Level 1 annotations as prior knowledge, we fine-tuned scANVI^21^, a tool that proved to be one of the top-performing integration methods^22^, to remove the batch effects between different studies (Fig. 2b). Under each of these major cell types, we further performed un-supervised graph-based clustering for in-depth annotations, which generated 17 stable cell types (Level 2) (Fig.2c).

**Fig. 2.**
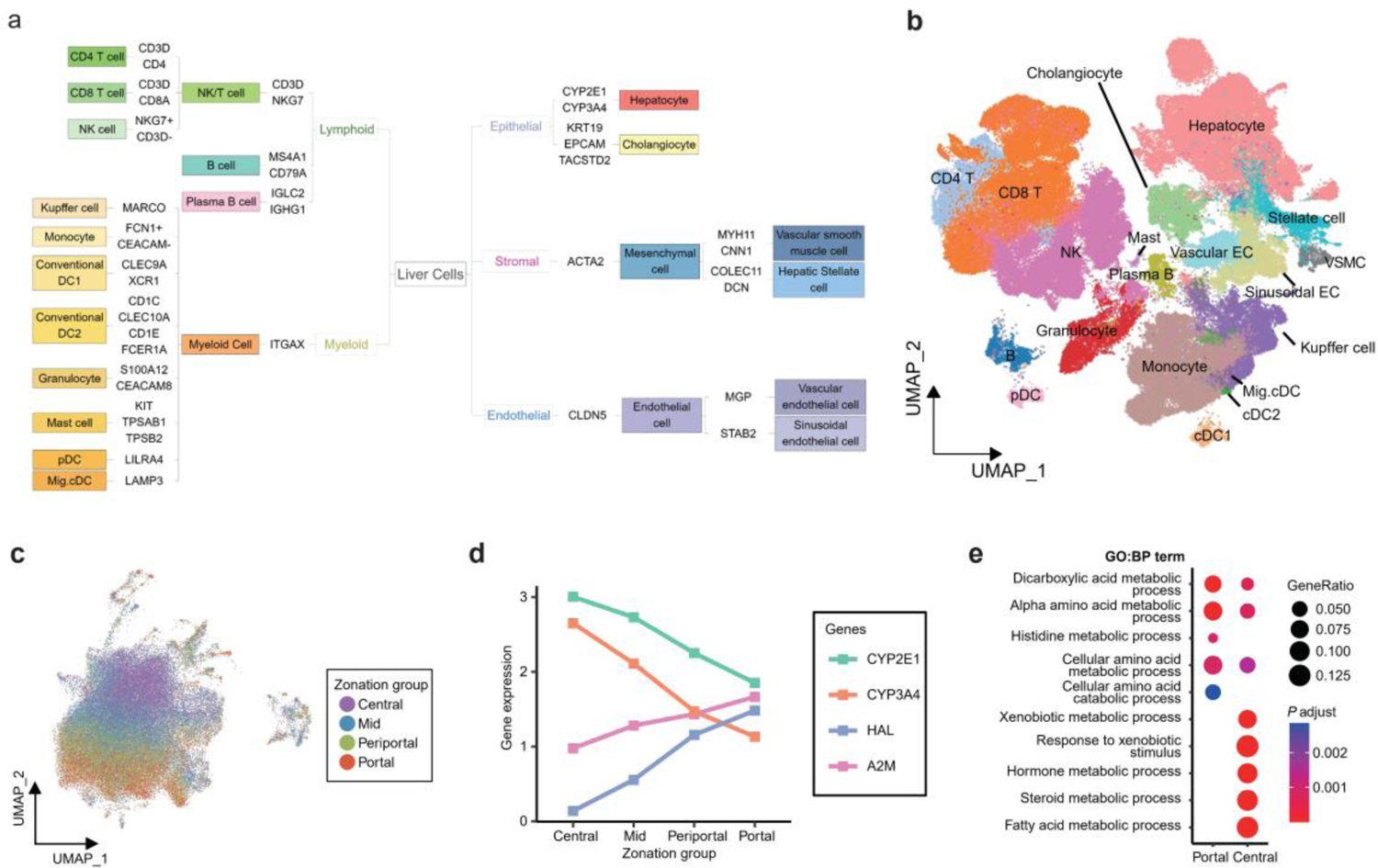
Construction of the normal reference map. **a**, The Level 1 and Level 2 uHAF tree of liver. b, The UMAP visualization of the normal reference map. c, The UMAP of hepatocytes (colors are zonation annotations). d, Gene expression of canonical markers *CYP2E1*, *CYP3A4*, *HAL* and *SBDS* across the CV-PV axis. e, The significantly enriched biological process (GO:BP terms with Benjamini–Hochberg-adjusted P < 0.05) of the genes upregulated in portal and central regions, respectively.

The hepatic lobule is the basic unit for liver function, with a central vein (CV) in the middle and portal vein in the six corners (PV). The hepatocytes have different states and functions along the CV to PV axis. The lobule can be roughly divided in four regions: central, middle, portal and peri-portal zones^5^ (in a few studies the portal and peri-portal zones are combined as a single zone^23,24^). We annotated hepatocyte zonation utilizing the gene signatures obtained from spatial transcriptomes (Methods, Fig. 2d, Extended Data Fig. 2a-c, Supplementary Table 3). The annotated zonation patterns can be validated by the expressions of multiple canonical marker genes: *CYP2E1* and *CYP3A4* gradually decreased along the CV-PV axis while *HAL* and *A2M* exhibited an opposite trend, highly consistent with previous studies^5,25^ (Fig. 2e). Functional enrichment analysis shows that the up-regulated genes of the hepatocytes annotated as in central region were enriched in xenobiotic metabolism pathways and the result of portal region was enriched in amino acid processing (Fig. 2f, Extended Data Fig. 2d), which was also consistent with the well-established cellular functions of the hepatocytes in different zones^26^.

This large scale of data collection enables us to primarily investigate the impact of population variables (e.g. gender and age) on the cell type compositions and subtle gene expression variations in a same cell type. We found no significant difference in the proportion of cell types between genders (Extended Data Fig. 3a, b). As the age increases, the proportion of different cell types underwent complex changes (Extended Data Fig. 3c, d). The effects of these variables on gene expression variations were also explored using generalized linear mixed models^27–29^. We found that the biological processes, including cellular response to ions (*MT1A* and *MT1E*), material transport (for example, cholesterol and sterol transport including *APOC2* and *APOM*), homeostasis maintenance (such as cholesterol, sterol, and lipid homeostasis including *AKR1C1*, *APOC2*, and *APOM*) and immune response (*C1A*, *CFHR2*, *RARRES2*) were significantly downregulated in monocytes within the elder population (Extended Data Fig. 3e).

### Deviated state analysis identifies diverse disease-associated cellular states

Compared to the normal liver, the cells in disease often exhibit certain cellular states “deviation”, which provide potential targets for treatment. To elucidate the state deviation extent of the cells under disease conditions, we developed LiverCT with a supervised and hierarchical ensemble learning framework to calculate a quantitative deviated score based on the normal reference map. The performance of LiverCT was firstly validated on predicting the cell type labels in the normal reference map (Methods, Extended Data Fig. 4a).

The collected disease datasets were mapped to the normal reference by LiverCT (Fig. 3a). We observed higher deviated scores in the cells from tumor (T) tissues compared to adjacent non-tumors (NT) in most cell types. In primary tumors, hepatocytes, cholangiocyte and granulocytes changed significantly, indicating strong deviation to the normal reference (Fig. 3b, Extended Data Fig. 4b). Hepatocytes and cholangiocytes were parenchymal cells that were prone to oncogenic transformations, making them distinct from normal tissues. A recent study also reported that neutrophils (major cell group of granulocytes) in liver cancer tissues exhibited significant gene expression changes compared to non-tumor tissues^16^.

**Fig. 3.**
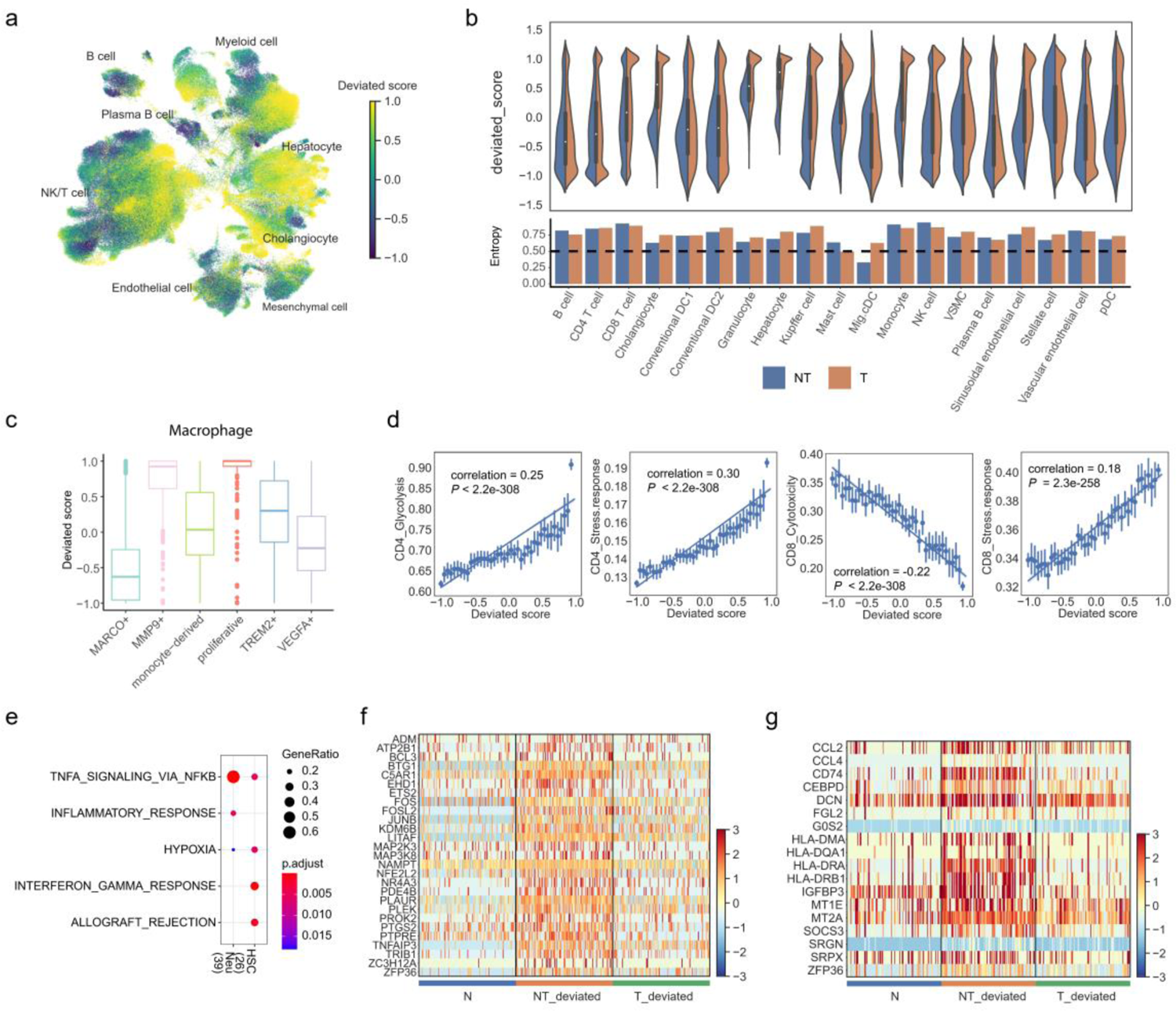
Deviated state analysis reveals the changes in adjacent non-tumor and tumor samples. a, The UMAP of the disease datasets with deviated scores. b. The deviated score distribution of each cell type at Level 2 (upper panel) and the entropy of the deviated state cells (below panel). c, Deviated scores of the macrophages with their original labels in Lu *et al*. datasets. d, Regplot of the deviated scores and T cell functional signatures for both CD4 and CD8 T cells. Pearson correlations were used to assess the associations. E, Enriched cancer hallmarks in the genes consistently upregulated in “deviated” neutrophils (Neu) and hepatic stellate cells (HSC) in adjacent non-tumor samples (adjusted P-value < 0.05). f,g, Heatmap of the expressions of the upregulated genes in the enriched hallmarks (Neu, f; HSC, g).

Furthermore, we tested the correctness of the deviated states on a HCC dataset published recently by Lu *et al*.^9^. They found the MMP9+ macrophages to be tumor-associated macrophages. It was observed that these MMP9+ macrophages had high deviated scores, with almost all of the cells scored positive (Fig. 3c). Also, there was no proliferative macrophage observed in the normal reference and this group got the highest deviated score in the HCC dataset. Besides, other known tumor-associated cell types such as regulatory T cells, pro-metastatic hepatocytes can also be successfully found with high deviated scores (Extended Data Fig. 4c). Taken together, above results showed that LiverCT can accurately retrieve the deviated state cells.

Tumor-infiltrating T cells have paved a novel way for tumor therapy^30^. We investigated the correlation between the function of T cells and their deviated scores^31^. The deviated scores of CD4 T cells have a positive correlation with the signatures of glycolysis and stress response. The scores of CD8 T cells show also a positive correlation with stress response signature, but a negative correlation with cytotoxicity (Supplementary Table 4, Fig. 3d). A recent pan-cancer study observed a strong association between the stress response and immunosuppression^31^, suggesting that the deviated score of T cells is a possible alternative indicator for immunotherapy response.

The adjacent non-tumor tissue (NT) presents a unique state between the normal and tumor^32^ and may provide additional information of the oncogenic transformation and recurrence^33^. Based on the cell type mapping results by LiverCT, we observed that the granulocytes, hepatocytes, and stellate cells had the highest number of the cells in deviated states. To unveil the NT’s unique expression features, we conducted differential expression analyses and focused on the consistently up-regulated genes in NT’s deviated cells. The up-regulated genes in both granulocytes and stellate cells were significantly enriched in the TNF-α signaling pathway (Fig. 3e-g, Supplementary Table 5), consistent with a pan-cancer study that observed NT-specific TNF-α signaling pathway activation^32^. Collectively, the deviated state analysis identified diverse disease-associated cellular states and illustrated the most susceptible cell types in disease.

### Intermediate state analysis reveals a population of tumor cells associated with poor prognosis in HCC

Intermediate or transition cellular states are widely induced in tumors. For instance, in human melanoma^34^, a transitional CD8 state is observed, and in peripheral neuroblastic tumors, an intermediate state is observed between adrenergic and mesenchymal neuroblasts^35^. To find the cells with intermediate states, LiverCT can also calculated an “intermediate” score for query cells (Fig. 4a). In the collected liver cancer datasets, cells between CD4-CD8, Mono-Macro, HSC-VSMC and LSEC-VEC exhibited high intermediate states ratio (Fig. 4b).

**Fig. 4.**
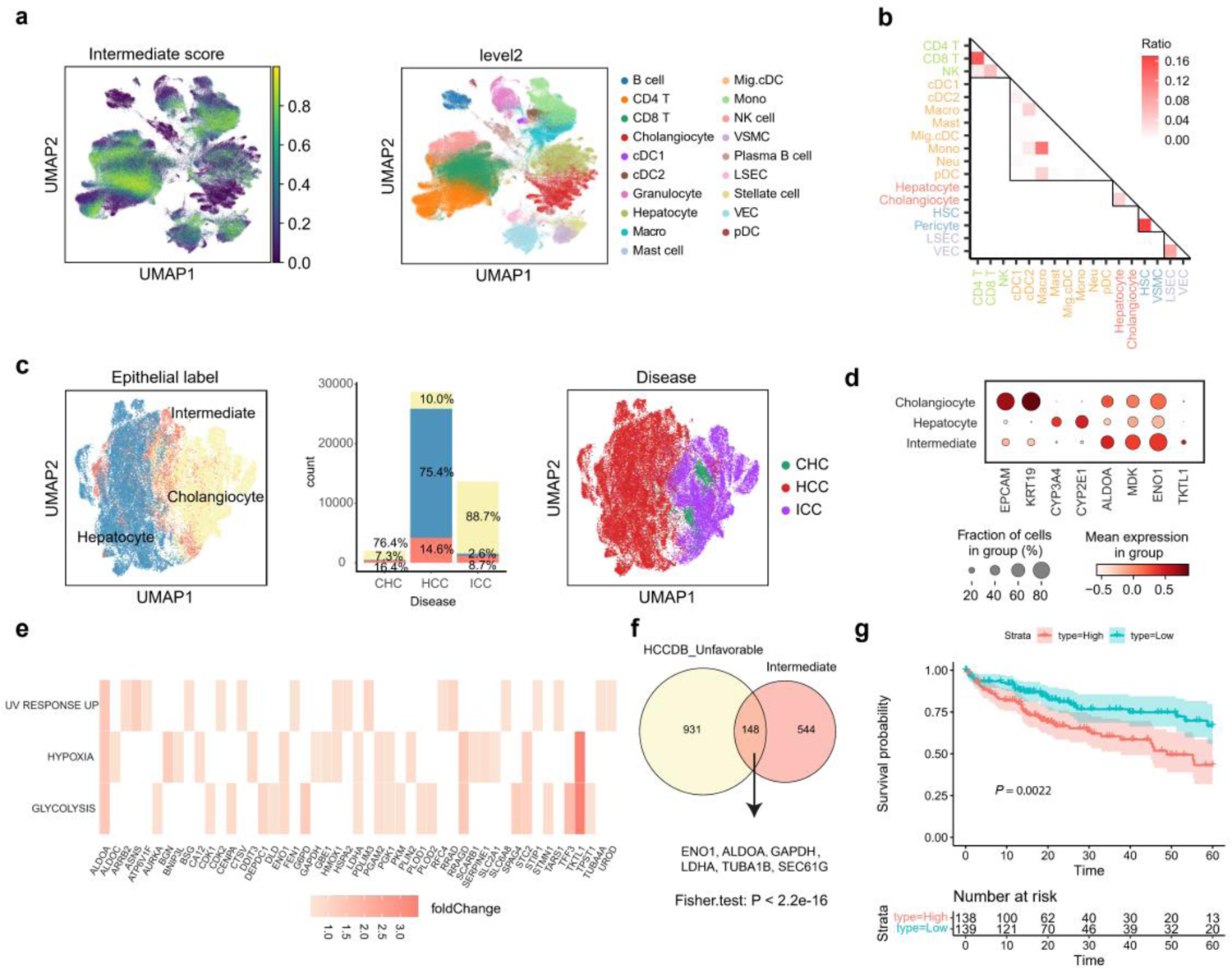
Intermediate state analysis enable the identification of the tumor cells associated poor prognosis. a, The UMAP visualization of the intermediate scores in the tumor samples. b, The ratio of the intermediate state cells occupying the proportions of the two “terminal” cell types. c, The UMAP visualization of the tumor (epithelial) cells annotated as hepatocyte-like, cholongiocyte-like and intermediate state cells by LiverCT. d, The expressions of selected marker genes for cholangiocytes and hepatocytes, and also the highly expressed genes in the intermediate state tumor cells. e, Three cancer hallmarks enriched in the upregulated genes in the intermediate state tumor cells (adjusted P < 0.05). f, Venn diagram of the genes upregulated in the intermediate state tumor cells and the unfavorable genes listed in HCCDB. g, The survival analysis of the TCGA HCC cohort based on the gene signature derived from the tumor cells with intermediate states (P value was calculated using log-rank test).

**Fig. 5.**
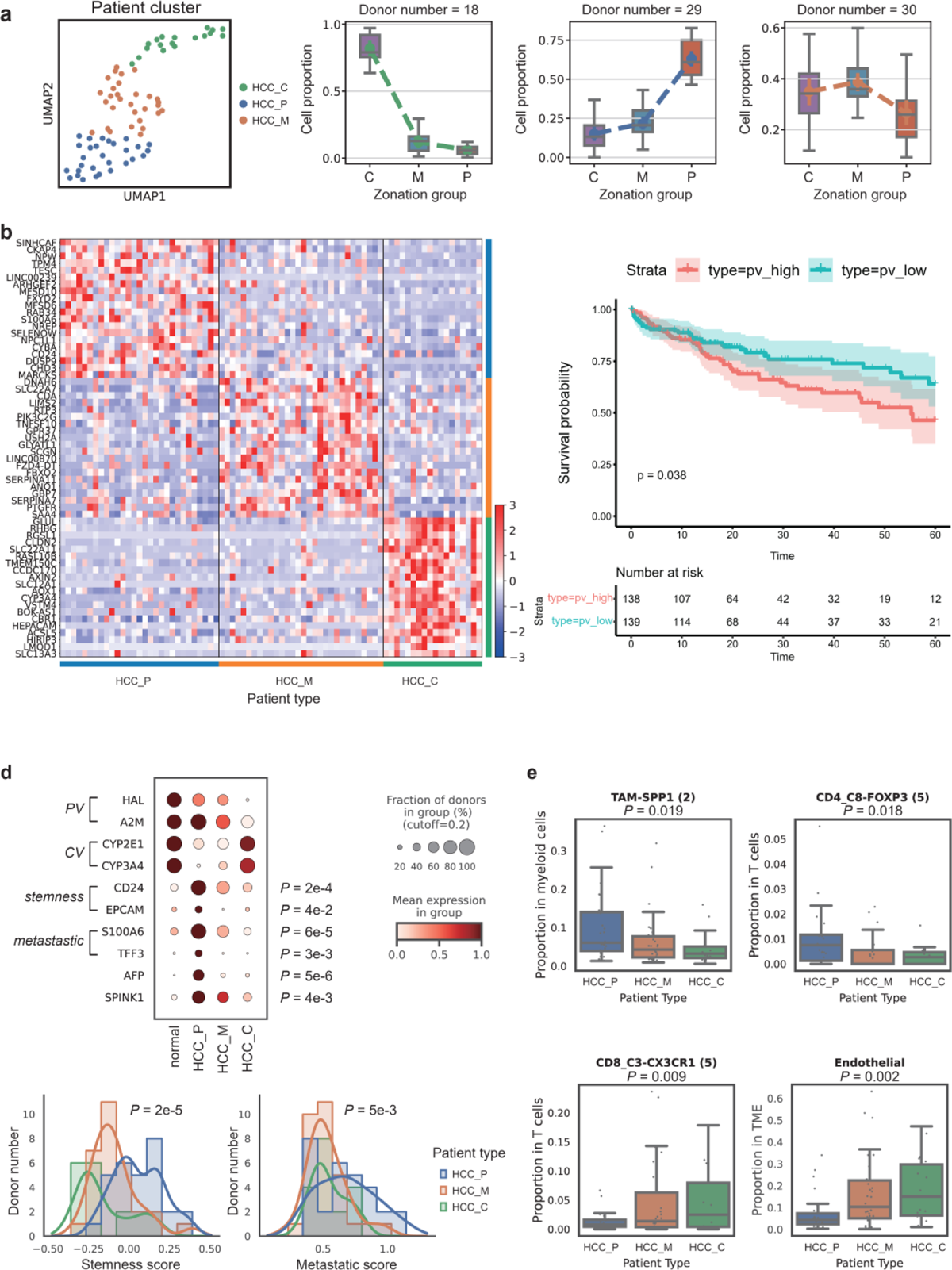
The discovery of HCC subtypes by tumor cell zonation mapping. a, The UMAP of the three HCC subtypes defined based on the distributions of LiverCT mapped tumor cells’ zonation states. b, The heatmap the differentially expressed genes among the three subtypes using the patient-level pseudo-bulk data. c, The survival analysis of the TCGA HCC patients based on the HCC_P pseudo-bulk signature (the P value was calculated by log-rank test). d, The expression of marker genes for central, peri-portal areas and HCC_P patients (top). The stemness score and metastatic score distribution among different patient types (bottom). e, The proportions of the TME cell subtypes in the three HCC subtypes.

Also, we observed that many tumor cells have high intermediate scores between the hepatocyte-cholangiocyte pair. Liver cancers, including HCC, ICC, and CHC, exhibited significant heterogeneity and can arise from diverse origins^36^. It has been reported that ICC could potentially arise from biliary-like cells that undergo trans-differentiation from hepatocytes, as well as from hepatic progenitor cells in addition to mature cholangiocytes^37^. So, we then tend to study the characteristics of those tumor cells with intermediate states.

LiverCT classified the malignant epithelial cells from tumor samples into hepatocyte, cholangiocyte and intermediate states (Methods, Fig. 4c). Notably, of the three types of liver cancer, the ratio of intermediate states in CHC was the highest, followed by HCC. Cholangiocytes exhibited high expression of *EPCAM* and *KAT19*, while hepatocytes expressed *CYP3A4* and *CYP2E1*. The intermediate states, however, under-expressed both these cell type markers. (Fig. 4d, Extended Data Fig. 5a).

To investigate the cellular features of the intermediate states, we performed a differential expression analysis, comparing the gene expressions of the tumor cells with intermediate states to the other tumor cells. We found a set of up-regulated genes associated with cell growth and development (*ALDOA*, *MDK*, *ENO1*, *TKTL1*) (Fig. 4e, Extended Data Fig. 5b, Supplementary Table 6). Among them, *ALDOA* was proved to serve as a driver for HCC cell growth under hypoxia^38^. *MDK* has been proposed as a multifunctional protein in HCC development, progression, metastasis, and recurrence^39^. The up-regulated genes were also enriched in glycolysis, hypoxia, and UV_ response_up besides cycling-related pathways (Fig. 4e, Extended Data Fig. 5c).

Subsequently, we explored the association between the intermediate states and prognosis in HCC using another database HCCDB which integrates multiple large-scale clinical cohorts to examine the gene expression variations in HCC^40^. Using Fisher’s exact test, we compared the poor prognosis-associated genes listed in HCCDB with the genes upregulated in the intermediate state tumor cells. Remarkably, the results demonstrated a significant overlap between these two gene lists, indicating a potential association between the intermediate states and the poor prognosis in HCC (Fig. 4f). We further examined the clinical relevance of the differentially expressed genes in intermediate states in the TCGA cohort^41^ and found that higher intermediate gene signature score is significantly associated with worse overall survival (Fig. 4g). Taken together, the intermediate states analysis revealed a group of tumor cells with multiple malignant features which located between the hepatocyte and cholangiocyte states.

Finally, we studied the relationship between the deviated scores and the intermediate scores. It was observed that MACRO+ macrophages (kuppfer cells) had low intermediate scores and deviated scores, consistent with its preference in non-tumor tissues. And while, MMP9+ macrophages get high deviated scores and low intermediate scores, which were also consistent with that MMP9+ macrophages were in a terminal state in HCC^9^ (Extended Data Fig. 5d).

### Tumor cell zonation tendency mapping defines novel HCC subtypes

Hepatocytes in different lobular zones display spatial-ordered functional heterogeneities. However, in HCC tumor tissues, the lobule-like patterns are usually lost. It is an interesting question that whether the malignantly transformed hepatocytes (tumor cells) still have zonation tendency and whether the tendency patterns are associated with clinical outcome.

Leveraging the second module of LiverCT, the HCC tumor cells were mapped into three zones as the P (Periportal/Portal) state, the M (Mid) state or the C (Central) state (Methods). We observed different distributions of tumor cells’ zonation labels in different patients. Based on the different compositions of the LiverCT annotated zonation labels of tumor cells, we divided the patients into three groups, namely HCC_P (dominated by “Periportal+Portal” state cells), HCC_M (dominated by “Mid” state cells), and HCC_C (dominated by “Central” state cells) (Fig. 6a) (Methods).

**Fig. 6.**
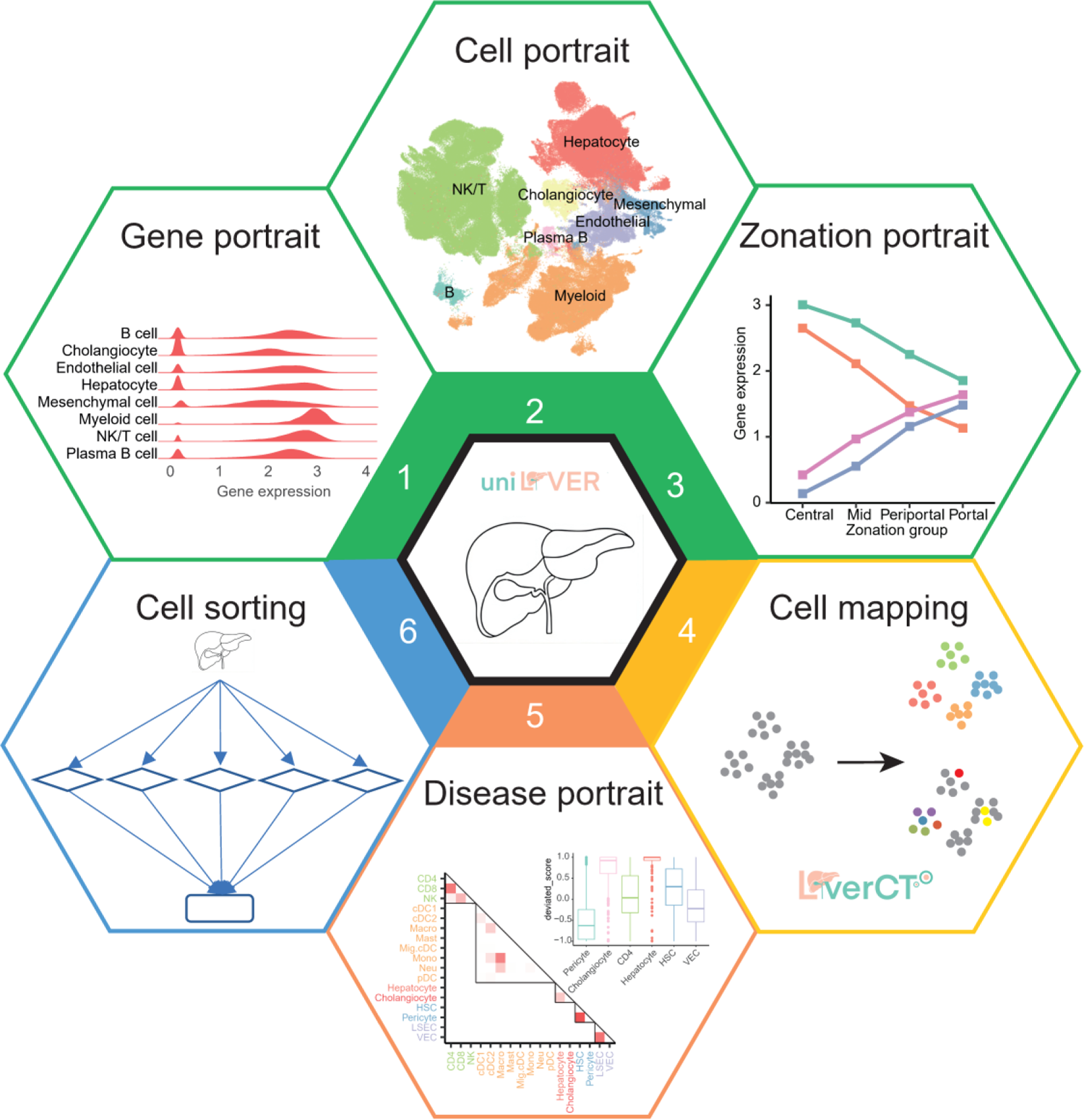
The database content and online tools of uniLIVER. The database consists of four portraits and two tools: (1) the *gene portrait* showing the gene expression among different cell types and subtypes within a lineage; (2) the *gell portrait* displaying uHAF tree and the cell-cell interaction within normal data; (3) the *zonation portrait* showing the gene expression within four zones in liver lobules; (4) the *cell mapping* page displaying the cell annotation and variant state identification as well as zonation reconstruction pipeline of LiverCT; (5) the *disease portrait* showing the characteristics of deviated states and intermediated states; (6) The *cell sorting* providing a one line tool which allows users to download data in uniLIVER flexibly.

To explore the distinctive features of the three HCC subtypes, we calculated the differentially expressed genes (DEGs) of each of the subtypes by using patient-specific pseudo-bulk data (Fig. 6b, Supplementary Table 7). Results show that in addition to several well-known zonal marker genes, different subtypes of HCC patients exhibited unique expression of many other non-zonal genes. Notably, *CD24* was among the top DEGs of the HCC_P subtype, suggesting a more malignant phenotype^42^. We further performed a survival analysis in TCGA^41^ bulk data, and found that higher HCC_P signature scores were significantly associated with poorer clinical prognosis (Fig. 6c). Multiple independent cohorts in HCCDB^40,43^ showed the similar observations (Extended Data Fig. 6a).

By scoring the curated stemness gene sets (*ANPEP*, *CD24*, *CD44*, *PROM1*, *EPCAM*)^44^ and metastatic gene sets^9^ on the pseudo-bulk data, we found statistically significant differences in the distribution of the stemness scores (*P* = 2e-5, ANOVA test) and the metastasis scores (*P* = 5e-3, ANOVA test) among the three HCC subtypes. The HCC_P patients showed much higher scores of cancer stemness and metastasis than the other two subtypes (Fig. 6c). Besides, in tumor cells of HCC_P, we observed the higher expression of *AFP*, which is a marker for fetal liver and a well-known marker for primary liver cancer. Additionally, *SPINK1*, a marker for hepatoblasts^45^ and a tumor-promoting factor^46^, exhibited up-regulation in tumor cells of HCC_P (Fig. 6c). These differences were not observed among the cells in different zones in normal liver, suggesting that these alterations should be associated with pathological changes (Extended Data Fig. 6b, c). Based on these findings, we speculated that the HCC_P subtype may correspond to a more poorly differentiated HCC phenotype.

Then, we investigated the dys-regulated metabolic processes in these HCC subtypes. Pseudo-bulk of HCC patients and normal donors were scored using a set of curated metabolic gene modules^47^. We found that urea cycle, which is normally conducted in portal and periportal regions^48^, is most significantly down-regulated in HCC_P patients. The finding suggested that tumor hepatocytes of HCC_P subtype presented a higher degree of urea cycle disorder (UCD), which correlated with poor prognosis^47^. Conversely, glycolysis was not a normal functional process of portal and periportal hepatocytes^23,49^, but we observed a slight upregulation of the glycolytic modules in tumor cells of HCC_P patients (Extended Data Fig. 6d).

We further conducted a detailed analysis of the characteristics of the tumor microenvironment (TME) in patients with HCC_P subtype. Utilizing scCancer2^50^, we transferred TME cell labels from multiple public datasets^15,51,52^ to our collected data. Our analysis revealed that HCC_P patients had a higher abundance of SPP1+ tumor-associated macrophages (TAMs) and FOXP3+ CD4 T cells compared to the other subtypes. In contrast, the proportions of CX3CR1+ CD8 T cells and endothelial cells were lower in HCC_P patients (Fig. 6e, Extended Data Fig. 6e). We noticed that SPP1+ TAMs were described as a potentially pro-tumorigenic/pro-metastatic subtype in colorectal cancer. Intriguingly, we found that cells annotated as SPP1+ TAMs in HCC TME displayed high expression levels of markers for MMP9+ macrophages^9^ (Extended Data Fig. 6f), which had been previously implicated in promoting HCC progression. These observations suggested that patients classified as HCC_P subtype might exhibit a relatively immunosuppressive TME.

Taken together, the zonation mapping of tumor cells by LiverCT defines novel HCC subtypes. Notably, HCC_P patients exhibited the worst overall survivals, characterized by increased expression of stemness and metastatic factors, along with the presence of metabolism dysregulation and an immunosuppressive microenvironment.

### A web-based portal of multidimensional portraits of the atlas

To facilitate a convenient browsing, we design a web-based portal for the atlas. It contains two user-friendly tools, namely *cell mapping* and *cell sorting*, and four portraits which are the *gene portrait*, *cell portrait*, *zonation portrait* and *disease portrait*, each portrait contains multiple different views (Fig. 6).

Detailed information of genes, cells and zonation of normal reference cell map can be found in the corresponding portraits. The *gene portrait* provides the expression distribution of the selected gene across cell types. The *cell portrait* portrays the uHAF tree of uniLIVER as well as the features of the selected cell type quantitively, including its number and cell-cell interaction. The *zonation portrait* provides the expression distribution of the selected gene across zonation and highly expressed genes (HEGs) of each zone.

For mapping new datasets onto the normal reference map, we developed a *cell mapping* tool LiverCT (freely available via uniLIVER website). The method mainly contains three parts, namely cell type classification, “variant” state identification, hepatocyte zonation reconstruction. When users input single-cell sequencing data, LiverCT will provide predicted cell types at Level 1 and Level 2. Additionally, it will provide two scores: the deviated score and the intermediate score, as well as cells the recommended thresholds for identifying the cells with deviated states or intermediate states, respectively.

Using LiverCT, we comprehensively annotated the disease data and constructed the *disease portrait*. The *disease portrait* shows the characteristics of the disease from two views: (1) *molecular view*; (2) *cellular view*. The *molecular view* presents the features of disease states and intermediate states, as well as the characteristics of disease cell types. The *cellular view* contains the statistical representation of cell type number and cell-cell interaction in the selected liver condition and difference compared with another liver condition.

In addition to the cell mapping tool LiverCT, the portal also embedded a *cell sorting* tool which allows users to download data in uniLIVER flexibly^53^. It is implemented on the hECA interactive web interface where users just need to input the filtering conditions to quickly obtain the desired data online and no longer need to download all the data and then filter it. This greatly saves time and is essential in the era of increasing data.

## Discussion

In this study, analogy to the genome sequence mapping, we have provided a machine learning based framework for disease “variant” analysis. As a tool for uniLIVER, LiverCT get several interesting findings by mapping disease datasets to the normal reference map. It finds that neutrophils and hepatic stellate cells are strongly deviated in adjacent tumor, and the intermediate-state tumor cells are associated with unfavorable outcomes in HCC.

The function of hepatocytes along the lobule radial axis is highly heterogeneous, which in turn results in differences of zonal patterns of drug responses and oncogenic transformation^48^. Although hepatocytes’ function is impaired by diseases, we posit that they still exhibit the characteristics of the CV-PV axis at the global transcriptome level. These characteristics might be influenced by both the microenvironment and long-term epigenetic phenomena^54,55^. Tumor cell zonation tendency mapping defines novel HCC subtypes. Among them, the HCC_P subtype has worst survival with a SPP1+ macrophage infiltrated suppressive immune microenvironment. Clinically, the HCC novel subtypes enable different therapy choices. Further investigation is needed to elucidate the molecular mechanisms by which tumor cells interacts with immune cells, ultimately resulting in a poorer prognosis in HCC_P patients.

Defining a comprehensive and refined normal reference map is essential but challenging, as it requires capturing both cellular and population variations^29^. LiverCT presents a promising opportunity to assess the saturation of the atlas. When incorporating new healthy datasets, we can evaluate whether any novel deviated states emerge. If no new cell types are discovered, we can consider the atlas to be ready.

However, if new cell types are detected, we can fine-tune the model until the identification of previously unobserved cellular states ceases. As data continues to accumulate, there is a recent surge in the development of large-scale models that have demonstrated state-of-the-art performance across a wide range of downstream tasks^56–59^. These models offer a promising opportunity to create a more comprehensive atlas. Also, with the development of spatial transcriptomics technology, it is now possible to further portray the spatial microenvironment of a cell, which is important to understand the cellular niches of liver.

## Methods

### Data collection and processing

In the current atlas, we archived 18 human liver datasets, including 6 healthy datasets, 1 cirrhosis dataset and 11 liver cancer datasets (Supplementary Table 1). For 17 publicly available datasets provided by dataset generators, we collected the expression matrix and processed it using Seurat pipline^60^. Besides the public datasets, we generated ∼30K healthy data and use scCancer^61^ pipeline to do quality control. The gene symbols were unified to the list of 43,878 HUGO Gene Nomenclature Committee (HGNC) approved symbols with the toolkit in hECA^53^, with withdrawn and alias symbols converted into HGNC approved symbols.

In addition, we collected phenotype information at multiple levels including donor, sample and cell. At the donor level, we gathered gender, age and fibrotic status if available. At the sample level, we categorized the sample status according to its location, harmonizing it as normal (N), primary tumor (T), non-tumor (NT), the joint area between the tumor and adjacent normal tissues (PJ), hepatic lymph node (HLN), metastatic lymph node (MLN), portal vein tumor thrombus (PVTT), Ascites (ASC), Blood (BLO) (Supplementary Table 1). At the cell level, we collected the original annotations and standardized them to the cell type at level 1 in the uHAF tree (Supplementary Table 2).

### Normal data integration and annotation

To visualize the cells in the normal reference map, the neighbor graph was built based on the 30 latent dimensions that were obtained from the scANVI output with the default parameter setting of sc.pp.neighbors function. The dimensionality of cells was further reduced using Uniform Manifold Approximation and Projection (UMAP) with sc.tl.umap function based on the neighbor graph built above. To determine the Level 1 label of cells, we used two methods. If the original study provided labels of cells, we would map those labels to the uHAF to obtain the Level 1 label. If not, marker genes would be used to identify the cell types.

To further annotate cells of each resulting Level 1 cluster, a new neighbor graph was built using 30 latent dimensions of scANVI. Clusters were classified into level 2 labels using marker genes.

### Normal hepatocyte zonation annotation

We annotated zonation labels for hepatocytes from the uniLIVER normal reference map. The zonation groups provided by Guilliams et al. (2022) for the human liver spatial transcriptome were used as the reference^5^.

For each Visium sample, we conducted Wilcoxon test to find differential expressed genes between C-spots (“Central”) and P-spots (“Periportal” +” Portal”). This step is implemented via rank_genes_groups() function in Scanpy^62^. In order to mitigate the impact of inter-individual variability, only genes showing significant zonal differences (pvals_adj < 0.01 for C-markers, and pvals_adj < 0.05 for P-markers) in more than 3 samples were considered. To accommodate scRNA-seq data characteristics, we filtered out genes with a mean log-normalized expression lower than 0.1 in hepatocytes from our single-cell data.

A min-max scaler is applied to each gene in the same sample first to preserving gradient information. Then, a spot’s score can be calculated as:

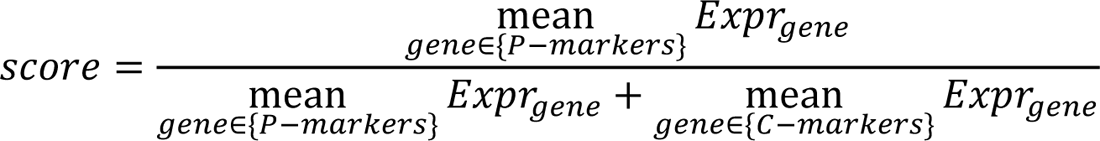

We visualized the original group labels and our defined score on Visium spots (Extended Data Fig. 2a). Furthermore, we plotted the score distribution for the four zonation labels (Extended Data Fig. 2b). These results demonstrated that the score can effectively indicate the location along the CV-PV axis in a healthy liver. This score can be applied equally to spots in spatial transcriptome, as well as cells in single-cell transcriptome.

For hepatocytes from uniLIVER normal reference map, we first conducted a quality control step. We removed non-viable cells with percentage of mitochondrial gene counts over 30%. Also, cells with an expressed gene number lower than 1000 were excluded from the subsequent annotation and analysis. These specific thresholds were determined based on the distribution of QC indicators obtained using Scanpy function calculate_qc_metrics() (Extended Data Fig. 2c).

To address distributional biases between spatial and single-cell transcriptomes, as well as variations in experimental techniques for single-cell sequencing, we conducted a correction step. Our hypothesis was that the scores for livers from healthy donors should exhibit a similar distribution. Therefore, we adjusted the mean and variance of the score distribution within each batch of single-cell transcriptome data to align with the corresponding distribution observed in the spatial transcriptome data.

We employed a normal function to fit the score distribution of each zonation label. The parameters were determined via maximum likelihood estimation: 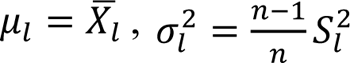. We transferred the previously fitted distribution to the single-cell data.

Bayesian estimation was utilized to infer the zonation group of each cell, assuming an equal prior probability for each zonation label:

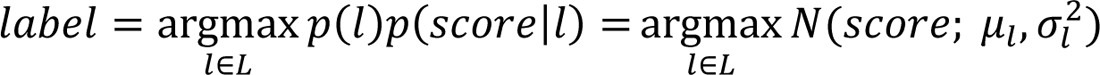

where L represents the set of the four zonation group labels (Central, Mid, Periportal, Portal).

### Modeling the effect of demographic covariates on gene programs

To model the effect of demographic covariates (gender and age) on gene programs, we performed the generalized linear mix model (GLMM). We first split cells by level 2 labels, then filtered out genes that were expressed in fewer than 10 cells. Sample-level pseudo-bulks, which were generated by summing gene counts across cells within each level 2 label for each sample, were used to fit the model. Pseudo-bulks were normalized using calcNormFactors function of edgeR with default parameter settings. Then voom^63^ was used to fit GLMM for differential expression and perform hypothesis test on fixed effects. Gene expression was modeled as:

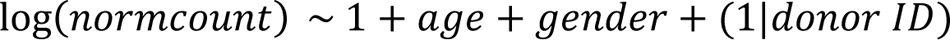

where the donor is treated as a random effect, and age and gender are modeled as fixed effects. We used the Benjamini-Hochberg procedure to correct the resulting p-values within each covariate. Significant genes (adjusted p-value < 0.05) were selected for gene set enrichment analysis using the enrichGO function in the clusterProfiler package^64^.

### LiverCT: a machine learning based cell-type mapping

We developed LiverCT (machine learning based Liver Cell Type mapping), to map new datasets onto the normal reference map. Two-level cell type labeling was provided by a hierarchical ensemble learning classifier. On the basis of accurate cell type prediction, LiverCT identified cells in “variant” states, which can be broadly categorized into two types: deviated states and intermediate states. Specifically, for hepatocytes, LiverCT further predicted zonal groups along the CV-PV axis at sub-lobule scale. The workflow of LiverCT is depicted in Extended Data Fig. 1b.

#### Batch correction

To mitigate batch effects between the query data and the normal reference, query datasets were projected to the common latent space of then normal reference map using scArches^20^, a transfer learning method. The parameter “encode_covariates” of the scANVI model was set to True to allow us to fine-tune the weights of newly introduced edges in the input layer. We conducted 20 epochs during the fine-tuning process of scArches. Subsequent models operated in this latent space.

#### Hierarchical ensemble learning cell type classification

The manually annotated labels served as the reference standard for classification. The process followed a hierarchical tree structure as shown in Fig. 2a, to improve the resolution of cell type labeling step by step. The query cells were first divided into 8 major cell types. Then within each major type, a finer-grained classification was carried out, resulting in 17 labels at the second level. Both layers of the classification were implemented using an ensemble learning model. It consisted of a Multi-layer Perceptron (MLP) classifier^65^, an XGBoost classifier^66^, a Logistic Regression classifier^67^ using one-vs-rest strategy and a Random Forest classifier^68^. A soft voting strategy was implemented to generate the predicted probabilities for each cell type. The algorithm was accelerated using parallel threading managed by the joblib (https://github.com/joblib/joblib) package.

#### Deviated states identification

We utilized a One-Class Support Vector Machine (OCSVM) for unsupervised novelty detection^69^. For each fine-grained label in the second level, a OCSVM model was trained. By delineating the contour of the feature space occupied by cells in the normal reference, the OCSVM model effectively identified cell states that deviated from the normal states. We first used a Radial Basis Function (RBF) Kernel to transform the initial observations to a non-linear feature space:

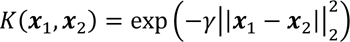

The feature map for the RBF kernel was approximated with the Nyestroem method for acceleration^70,71^, using sklearn.kernel_approximation.Nystroem(). Then, a linear OCSVM was performed in the transformed feature space. The OCSVM model was solved using Stochastic Gradient Descent (SGD). This algorithm was chosen due to its efficiency in processing large training sets. The optimization problem was defined as follows:

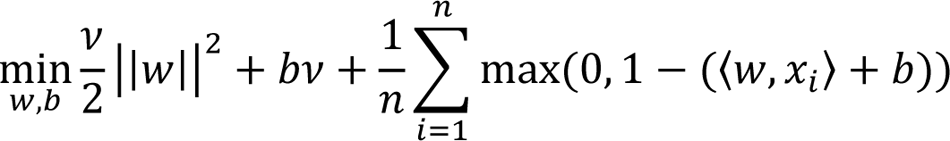

where *w* and *b* represented the linear coefficient and the intercept to be optimized, and *v* was a hyperparameter. The hyperparameters of the model were automatically set based on the distribution of the training data. Specifically, the parameter γ of the RBF kernel was set to 1/(*n_features ∗ var(X)*) as suggested by the sklearn library. The parameter *v* was incrementally increased until 10% of the training data was detected as outliers. Cells located outside the frontier-delimited subspace were annotated as under “deviated” states. Euclidean distances between query data observations to the frontier hypersphere were calculated. These distances were then normalized by the 80th percentile value for each model. Subsequently, the normalized distances were negated and truncated between −1 and 1, resulting in the deviated scores. Higher deviated scores represented larger deviations from the normal distribution.

#### Intermediate states identification

We assumed that intermediate states only existed between cell types under the same major type. A special case is hepatocytes and cholangiocytes, where an intermediate state between the two has been demonstrated to exist in certain disease conditions^36^.Therefore, even though they are already distinguished at the major cell type level, we have still identified the intermediate state between them. We employed a one-vs-one SVM model^72^ to identify the classification boundaries between the top two classes to which the cell was most likely to belong. The pipeline consisted of a standard scalar and a linear SVM optimized using SGD. We calculated Euclidean distances of the samples to the separating hyperplane, then used a generalized RBF kernel to transform distances to scores between 0 and 1:

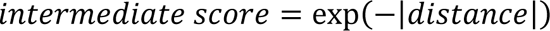

The higher intermediate score represented that the cell tended to be more intermediate between the two types. A threshold was determined as 0.6 manually to classify cells with scores above it as being in intermediate states.

#### Tumor hepatocyte zonation states mapping

In the context of disease data, particularly hepatocellular carcinoma (HCC), the expression of many zonal landmark genes was found to be absent or exhibited a loss of gradient. As a result, the scoring approach that relied on a subset of genes defined in a healthy liver was unsuitable for analyzing disease data. To overcome this, we employed a supervised learning classifier trained on our normal reference, and used it to transfer zonation labels to the disease data.

To address the individual heterogeneity of human hepatocytes, we used donor ID as batch labels to train a scANVI model. This model generated a 30-dimensional latent space with batch corrected. For the input features of the scANVI model, we identified 2000 highly variable genes (HVGs). The selection of HVGs was performed using the “seurat_v3” flavor provided by the Scanpy pipeline.

We chose the Random Forest algorithm, which employs feature sampling steps to ensure reliable classifications even when features are missing. This attribute makes it particularly suitable for analyzing disease state data. We used the low-dimensional latent vectors as input for training and implemented the algorithm using sklearn.ensemble.RandomForestClassifier with 100 estimators.

We used manually annotated zonation groups as the reference standard and implemented a more general categorization approach by using three classification labels for the training process. Specifically, we combined the Periportal and Portal regions, leading to three labels: C for Central, M for Mid, and P for Periportal+Portal.

For disease data, we utilized the transfer learning method, scArches, to acquire latent space representations consistent with the reference. During the scArches surgery process, 20 epochs of fine-tuning were performed. We then used the trained Random Forest classifier to predict zonation label for each individual cell.

We built LiverCT on Python (3.9.7), using the following packages: numpy (1.22.4), scipy (1.8.1), pandas (1.4.3), anndata (0.8.0), scanpy (1.9.1), scArches (0.5.9), joblib (1.1.0), scikit_learn (1.1.1), xgboost (1.7.6). The code is open-sourced at Github (https://github.com/fyh18/LiverCT).

### Variant states analysis

We sampled up to 1000 cells from each donor to maintain the balance of patient cell number. After quality control, 272,464 cells were left to constitute the core disease data.

In the T cell function analysis section, we use “sc.tl.score_genes” in scanpy to add module score to each cell. Pearson correlation was employed to assess the correlation between T cell function and deviation scores.

The intermediate gene signature was derived by filtering genes based on specific criteria, including a log fold change (FC) greater than 0.5 and an adjusted p-value (pvals_adj) lower than 0.01.

### HCC Classification

We analyzed cells annotated as hepatocytes by LiverCT from the core disease data. Only samples from HCC primary tumors were included. Samples with less than 50 hepatocyte-like cells were filtered out. We then calculated the proportion of cells with the three zonal labels (“C”, “M” and “P”) for each patient, resulting in a *n* × 3 matrix where each row represented a donor and each column represented the cell proportion of a certain zone. We referred to this matrix as “zonal proportion space of patients”. Subsequentially, we performed a 3-cluster spectral clustering within this space to classify three HCC subtypes, namely HCC_C, HCC_M and HCC_P.

We summed up the counts of all HCC tumor hepatocytes for each patient, and then perform log-normalization to acquire pseudo bulk data.

### Portraits of uniLIVER

#### Gene portrait

We provided the expression distribution of the selected gene across cell types. The ridge plots showed the non-zero expression distributions in different cell types. The number at the right of the ridge plots showed the non-zero percentages of the expression values.

#### Cell portrait

CellChat^73^ was employed to infer cell-cell interactions by analyzing the expression patterns of known ligand-receptor pairs across diverse cell types. We followed the official workflow with default parameters.

#### Zonation portrait

We provided the average expression values of the selected gene across zonation. Besides, differentially expressed genes of different zones can be visualized by heatmap.

#### Disease portrait

In *Molecular view*, we present the features of disease states and intermediate states, as well as the characteristics of disease cell types. The former two are compared within a specific disease, while the latter is compared between different disease conditions. In deviated state section, deviated score distribution in level 2 is displayed and we can see the most susceptible cell type. By selecting a cell type, differentially expressed genes in deviated states compared with normal states are shown. Similarly, in *intermediate state* section, the ratio of intermediate states between two cell type is shown and we can see the differentially expressed genes in intermediate states compared with the other two cell types. The *Disease cell type* section displays differentially expressed genes (DEGs) between the selected disease and another condition in the same cell type. Enrichment analysis is conducted based on the DEGs.

## Data availability

The uniLIVER website is publicly accessible via [https://liver.unifiedcellatlas.org]. The normal reference map and core disease data (processed as data matrix) are publicly available through the *databrowser* section and can be easily downloaded from *download* section in the web server. The source codes, trained models and documents of LiverCT are also provided at the website.

## Acknowledgements

This publication is part of the Human Cell Atlas – https://www.humancellatlas.org/publications/. We thank Qiuyu Lian, Qinglin Mei, Yiran Shan, Xinqi Li, Qifan Hu and Nan Yan and Yifan Sun for their help on this work. This work is funded by the National Key Research and Development Program of China (No. 2021YFF1200901) and the National Natural Science Foundation of China (Nos. 61721003, 62133006 and 92268104).

## Author contributions

J.G. conceived the study. Y.W., Y. F. and Y.M. collected datasets involved in the study. Y.W. and Y.F. designed the LiverCT algorithm. Y.F. implemented the LiverCT algorithm. Y.M. performed the unsupervised annotation experiment. Y.W. and Y.F. designed the biological applications. Y.M. imported data into the database. Y.F., Y.W., Y.L. and Y.M. designed the web page of uniLIVER. M.Y., R.Y., A.C. deployed the database. G.D., J.D. and Y.C. collected the clinical samples. Z.C., Y.C., W.L., W.G., J.D., X.Z. and Y.W. provided advice on experiments. Y.W., Y. F. and Y.M. wrote the manuscript. J.G. supervised the computational analysis. All authors agreed on the final version of the manuscript.

## Additional information

Extended Data Fig 1-6

Supplementary Table 1-7

## References

1. Hashimshony, T., Wagner, F., Sher, N. & Yanai, I. CEL-Seq: Single-Cell RNA-Seq by Multiplexed Linear Amplification. Cell Reports 2, 666–673 (2012).

2. Tang, F. et al. mRNA-Seq whole-transcriptome analysis of a single cell. Nature Methods 6, 377–382 (2009).

3. Aizarani, N. et al. A human liver cell atlas reveals heterogeneity and epithelial progenitors. Nature 572, 199–204 (2019).

4. Ramachandran, P. et al. Resolving the fibrotic niche of human liver cirrhosis at single-cell level. Nature 575, 512–518 (2019).

5. Guilliams, M. et al. Spatial proteogenomics reveals distinct and evolutionarily conserved hepatic macrophage niches. Cell 185, 379–396. e338 (2022).

6. Payen, V. L. et al. Single-cell RNA sequencing of human liver reveals hepatic stellate cell heterogeneity. JHEP Reports 3, 100278 (2021).

7. Andrews, T. S. et al. Single-cell, single-nucleus, and spatial RNA sequencing of the human liver identifies cholangiocyte and mesenchymal heterogeneity. Hepatology Communications 6, 821–840 (2022).

8. Losic, B. et al. Intratumoral heterogeneity and clonal evolution in liver cancer. Nature Communications 11, 291 (2020).

9. Lu, Y. et al. A single-cell atlas of the multicellular ecosystem of primary and metastatic hepatocellular carcinoma. Nature Communications 13, 4594 (2022).

10. Ma, L. et al. Single-cell atlas of tumor cell evolution in response to therapy in hepatocellular carcinoma and intrahepatic cholangiocarcinoma. Journal of hepatology 75, 1397–1408 (2021).

11. Massalha, H., et al. A single cell atlas of the human liver tumor microenvironment. Zenodo (2030).

12. Sun, Y. et al. Single-cell landscape of the ecosystem in early-relapse hepatocellular carcinoma. Cell 184, 404–421. e416 (2021).

13. Qi, Z. et al. Integrated multiomic analysis reveals comprehensive tumour heterogeneity and novel immunophenotypic classification in hepatocellular carcinomas. Gut 68, 2019 (2019).

14. Zhang, M. et al. Single-cell transcriptomic architecture and intercellular crosstalk of human intrahepatic cholangiocarcinoma. Journal of Hepatology 73, 1118–1130 (2020).

15. Zheng, C. et al. Landscape of Infiltrating T Cells in Liver Cancer Revealed by Single-Cell Sequencing. Cell 169, 1342–1356.e1316 (2017).

16. Xue, R. et al. Liver tumour immune microenvironment subtypes and neutrophil heterogeneity. Nature, 1–7 (2022).

17. Ma, L. et al. Multiregional single-cell dissection of tumor and immune cells reveals stable lock-and-key features in liver cancer. Nature Communications 13, 7533 (2022).

18. Liu, Y. et al. Identification of a tumour immune barrier in the HCC microenvironment that determines the efficacy of immunotherapy. Journal of Hepatology 78, 770–782 (2023).

19. Chen, S. et al. hECA: The cell-centric assembly of a cell atlas. iScience 25, 104318 (2022).

20. Lotfollahi, M. et al. Mapping single-cell data to reference atlases by transfer learning. Nature biotechnology 40, 121–130 (2022).

21. Gayoso, A. et al. A Python library for probabilistic analysis of single-cell omics data. Nature Biotechnology 40, 163–166 (2022).

22. Luecken, M. D. et al. Benchmarking atlas-level data integration in single-cell genomics. Nature Methods 19, 41–50 (2022).

23. Yang, Q., Zhang, S., Ma, J., Liu, S. & Chen, S. In Search of Zonation Markers to Identify Liver Functional Disorders. Oxidative Medicine and Cellular Longevity 2020, 9374896 (2020).

24. Paris, J. & Henderson, N. C. Liver zonation, revisited. Hepatology 76 (2022).

25. Halpern, K. B. et al. Single-cell spatial reconstruction reveals global division of labour in the mammalian liver. Nature 542, 352–356 (2017).

26. Sasse, D., Katz, N. & Jungermann, K. FUNCTIONAL HETEROGENEITY OF RAT-LIVER PARENCHYMA AND OF ISOLATED HEPATOCYTES. FEBS LETTERS 57, 83–88 (1975).

27. Squair, J. W. et al. Confronting false discoveries in single-cell differential expression. Nature Communications 12, 5692 (2021).

28. Crowell, H. L. et al. muscat detects subpopulation-specific state transitions from multi-sample multi-condition single-cell transcriptomics data. Nature Communications 11, 6077 (2020).

29. Sikkema, L. et al. An integrated cell atlas of the lung in health and disease. Nature Medicine 29, 1563–1577 (2023).

30. Zhang, Y. & Zhang, Z. The history and advances in cancer immunotherapy: understanding the characteristics of tumor-infiltrating immune cells and their therapeutic implications. Cellular & Molecular Immunology 17, 807–821 (2020).

31. Chu, Y. et al. Pan-cancer T cell atlas links a cellular stress response state to immunotherapy resistance. Nature Medicine 29, 1550–1562 (2023).

32. Aran, D. et al. Comprehensive analysis of normal adjacent to tumor transcriptomes. Nature Communications 8, 1077 (2017).

33. Kim, J. et al. Transcriptomes of the tumor-adjacent normal tissues are more informative than tumors in predicting recurrence in colorectal cancer patients. Journal of Translational Medicine 21, 209 (2023).

34. Li, H. et al. Dysfunctional CD8 T Cells Form a Proliferative, Dynamically Regulated Compartment within Human Melanoma. Cell 176, 775–789.e718 (2019).

35. Yuan, X. et al. Single-cell profiling of peripheral neuroblastic tumors identifies an aggressive transitional state that bridges an adrenergic-mesenchymal trajectory. Cell Reports 41, 111455 (2022).

36. Sia, D., Villanueva, A., Friedman, S. L. & Llovet, J. M. Liver Cancer Cell of Origin, Molecular Class, and Effects on Patient Prognosis. Gastroenterology 152, 745–761 (2017).

37. Fan, B. et al. Cholangiocarcinomas can originate from hepatocytes in mice. The Journal of Clinical Investigation 122, 2911–2915 (2012).

38. Niu, Y. et al. Loss-of-Function Genetic Screening Identifies Aldolase A as an Essential Driver for Liver Cancer Cell Growth Under Hypoxia. Hepatology 74 (2021).

39. Gowhari Shabgah, A., et al. Shedding more light on the role of Midkine in hepatocellular carcinoma: New perspectives on diagnosis and therapy. Iubmb Life 73, 659–669 (2021).

40. Lian, Q. et al. HCCDB: a database of hepatocellular carcinoma expression atlas. Genomics, proteomics & bioinformatics 16, 269–275 (2018).

41. Chang, K. et al. The Cancer Genome Atlas Pan-Cancer analysis project. Nature Genetics 45, 1113–1120 (2013).

42. Barkal, A. A. et al. CD24 signalling through macrophage Siglec-10 is a target for cancer immunotherapy. Nature 572, 392–396 (2019).

43. Ziming, J., et al. HCCDB v2.0: Decompose the Expression Variations by Single-cell RNA-seq and Spatial Transcriptomics in HCC. *bioRxiv*, 2023.2006.2015.545045 (2023).

44. Tsui, Y.-M., Chan, L.-K. & Ng, I. O.-L. Cancer stemness in hepatocellular carcinoma: mechanisms and translational potential. British Journal of Cancer 122, 1428–1440 (2020).

45. Wesley, B. T. et al. Single-cell atlas of human liver development reveals pathways directing hepatic cell fates. Nature Cell Biology 24, 1487–1498 (2022).

46. Chen, F. et al. Targeting SPINK1 in the damaged tumour microenvironment alleviates therapeutic resistance. Nature Communications 9, 4315 (2018).

47. Wu, T. et al. Discovery of a Carbamoyl Phosphate Synthetase 1–Deficient HCC Subtype With Therapeutic Potential Through Integrative Genomic and Experimental Analysis. Hepatology 74 (2021).

48. Ben-Moshe, S. & Itzkovitz, S. Spatial heterogeneity in the mammalian liver. Nature Reviews Gastroenterology & Hepatology 16, 395–410 (2019).

49. Manco, R. & Itzkovitz, S. Liver zonation. JOURNAL OF HEPATOLOGY 74, 466–468 (2021).

50. Zeyu, C. et al. scCancer2: data-driven in-depth annotations of the tumor microenvironment at single-level resolution. bioRxiv, 2023.2008.2022.554137 (2023).

51. Guo, X. et al. Global characterization of T cells in non-small-cell lung cancer by single-cell sequencing. Nature Medicine 24, 978–985 (2018).

52. Zhang, L. et al. Single-Cell Analyses Inform Mechanisms of Myeloid-Targeted Therapies in Colon Cancer. Cell 181, 442–459.e429 (2020).

53. Chen, S. et al. hECA: The cell-centric assembly of a cell atlas. Iscience 25, 104318 (2022).

54. Brosch, M. et al. Epigenomic map of human liver reveals principles of zonated morphogenic and metabolic control. Nature Communications 9, 4150 (2018).

55. Ben-Moshe, S. et al. Spatial sorting enables comprehensive characterization of liver zonation. Nature Metabolism 1, 899–911 (2019).

56. Theodoris, C. V. et al. Transfer learning enables predictions in network biology. Nature 618, 616–624 (2023).

57. Haotian, C., Chloe, W., Hassaan, M. & Bo, W. scGPT: Towards Building a Foundation Model for Single-Cell Multi-omics Using Generative AI. *bioRxiv*, 2023.2004.2030.538439 (2023).

58. Yang, F. et al. scBERT as a large-scale pretrained deep language model for cell type annotation of single-cell RNA-seq data. Nature Machine Intelligence 4, 852–866 (2022).

59. Minsheng, H. et al. Large Scale Foundation Model on Single-cell Transcriptomics. bioRxiv, 2023.2005.2029.542705 (2023).

60. Stuart, T. et al. Comprehensive integration of single-cell data. Cell 177, 1888–1902. e1821 (2019).

61. Guo, W. et al. scCancer: a package for automated processing of single-cell RNA-seq data in cancer. Briefings in Bioinformatics 22 (2020).

62. Wolf, F. A., Angerer, P. & Theis, F. J. SCANPY: large-scale single-cell gene expression data analysis. Genome Biology 19, 15 (2018).

63. Law, C. W., Chen, Y., Shi, W. & Smyth, G. K. voom: precision weights unlock linear model analysis tools for RNA-seq read counts. Genome Biology 15, R29 (2014).

64. Wu, T. et al. clusterProfiler 4.0: A universal enrichment tool for interpreting omics data. The Innovation 2, 100141 (2021).

65. Popescu, M.-C., Balas, V., Perescu-Popescu, L. & Mastorakis, N. Multilayer perceptron and neural networks. WSEAS Transactions on Circuits and Systems 8 (2009).

66. Chen, T. & Guestrin, C. in *Proceedings of the 22nd ACM SIGKDD International Conference on Knowledge Discovery and Data Mining* 785–794 (Association for Computing Machinery, San Francisco, California, USA, 2016).

67. Friedman, J., Hastie, T. & Tibshirani, R. Regularization Paths for Generalized Linear Models via Coordinate Descent. JOURNAL OF STATISTICAL SOFTWARE 33, 1–22 (2010).

68. Breiman, L. Random Forests. Machine Learning 45, 5–32 (2001).

69. Schölkopf, B., Platt, J. C., Shawe-Taylor, J., Smola, A. J. & Williamson, R. C. Estimating the Support of a High-Dimensional Distribution. Neural Computation 13, 1443–1471 (2001).

70. Williams, C. K. I. & Seeger, M. in Proceedings of the 13th International Conference on Neural Information Processing Systems 661–667 (MIT Press, Denver, CO, 2000).

71. Yang, T., Li, Y.-F., Mahdavi, M., Jin, R. & Zhou, Z.-H. in NIPS.

72. Cortes, C. & Vapnik, V. Support-vector networks. Machine Learning 20, 273–297 (1995).

73. Jin, S. et al. Inference and analysis of cell-cell communication using CellChat. Nature Communications 12, 1088 (2021).

