## Extended Data Fig 1-6 for "uniLIVER: a Human Liver Cell Atlas for Data-Driven Cellular State Mapping"

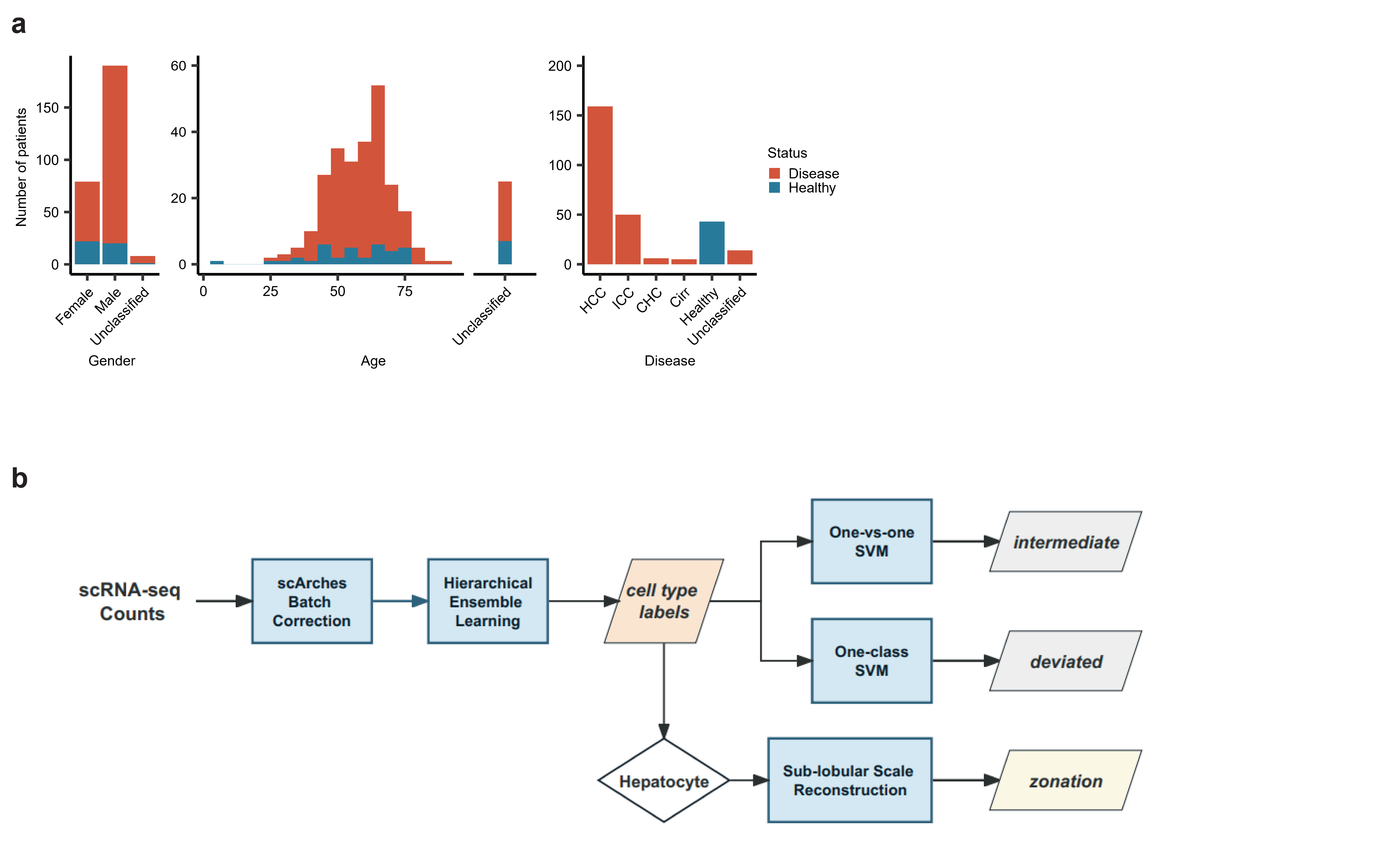

**Extended Data Fig. 1 | Summary of uniLIVER.** a, Gender, age and disease composition in uniLIVER. b, workflow of LiverCT including cell type classification, “Variant” state identification and hepatocyte zonation reconstruction.

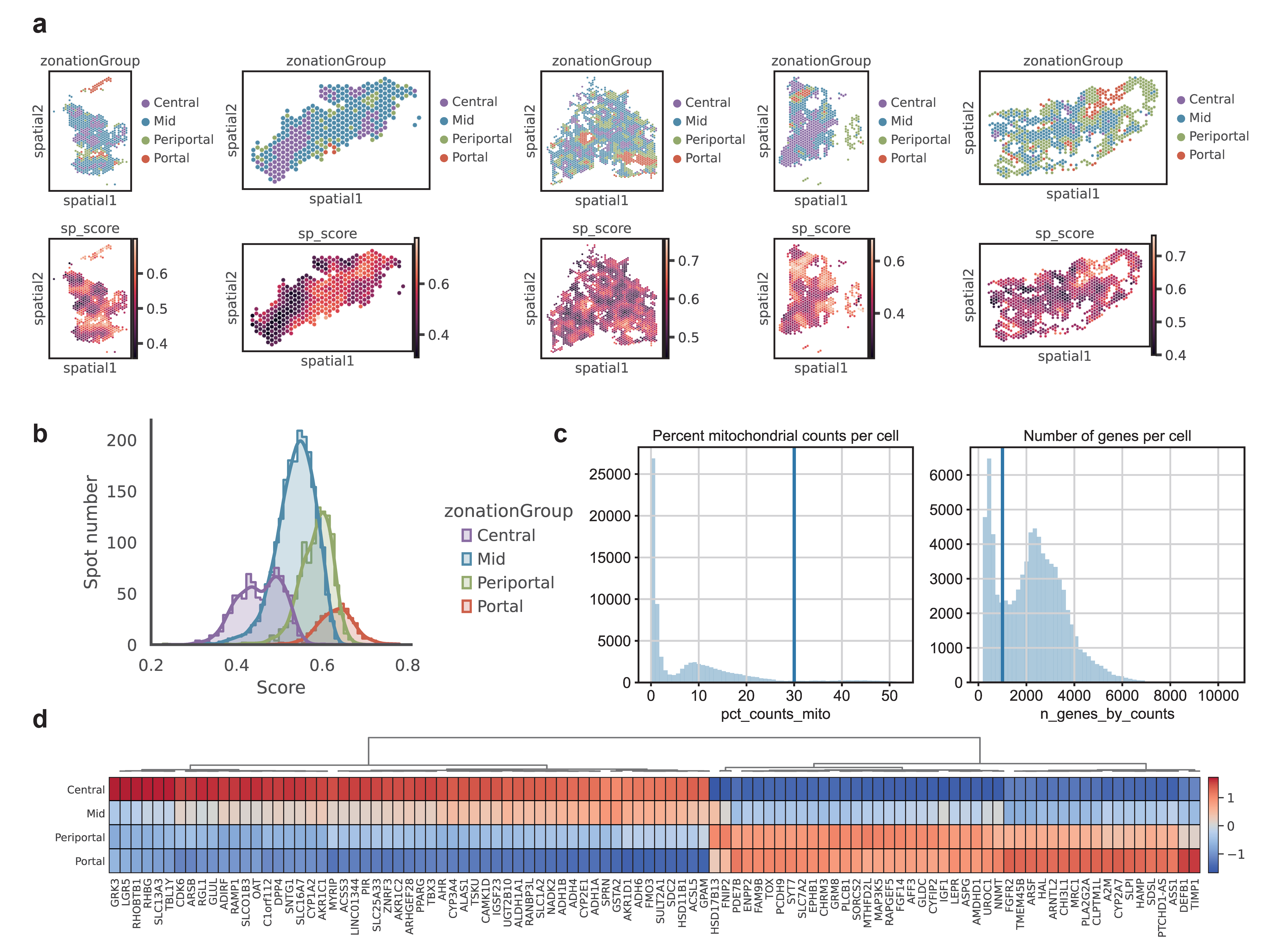

**Extended Data Fig. 2 | Normal hepatocyte zonation annotation. a,** zonation pattern labeled by Guilliams *et al.* (top) and spatial scores (bottom) mapped onto liver tissue. b, distribution of spatial scores across four zones. c, Distribution of percent mitochondial counts per cell in normal hepatocytes, the threshold for quality control is represented by a vertical line (left); Distribution of the number of genes per cell in normal hepatocytes, the threshold for quality control is represented by a vertical line (right). d, top DEGs for four zones.

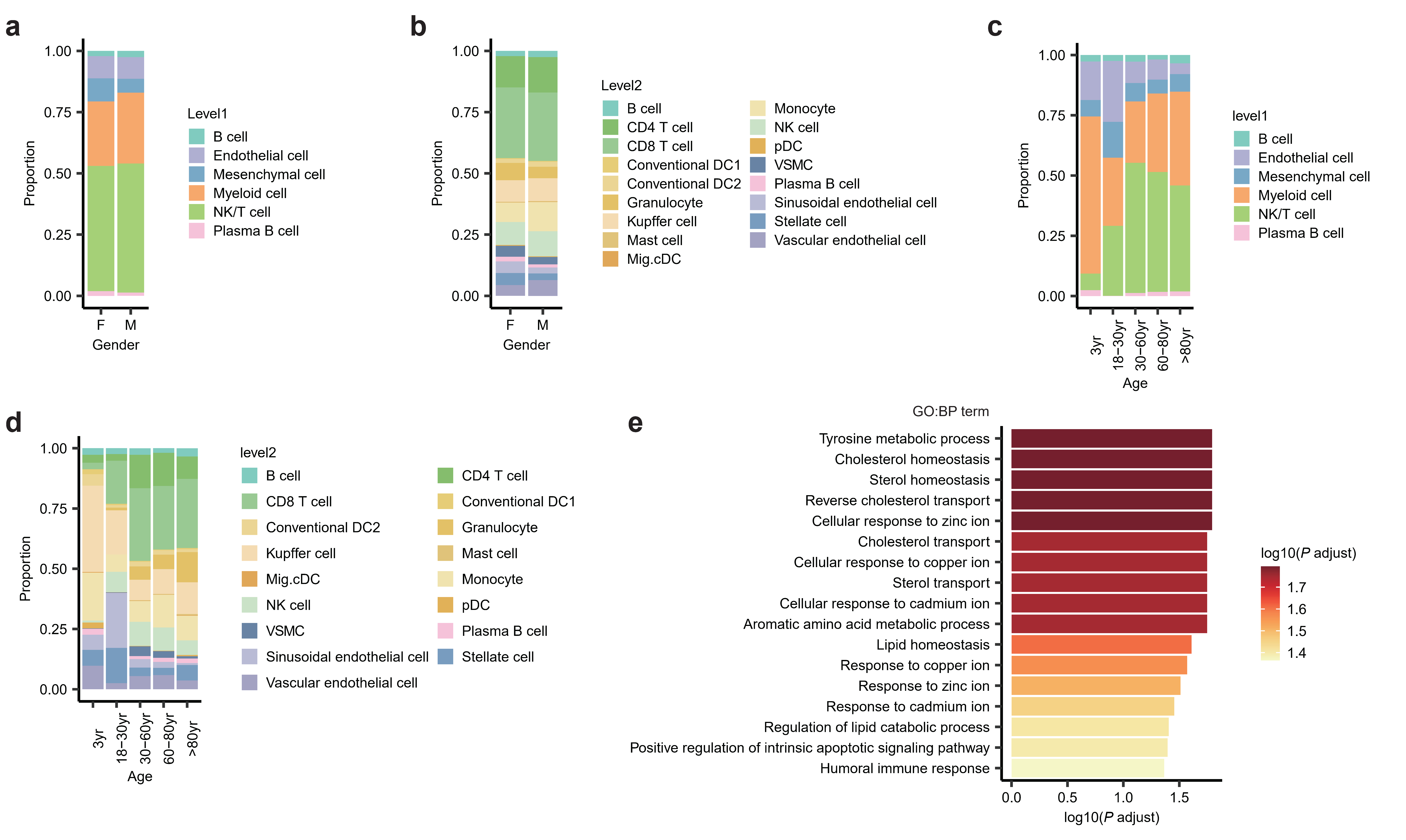

**Extended Data Fig. 3 | The impact of phenotypes on cell quantity and gene expression levels.** a, The relationship between gender and the proportion of Level 1 cell type labels in 10X data. b, The relationship between gender and the proportion of Level 2 cell type labels in 10X data. c, The relationship between age and the proportion of Level 1 cell type labels in 10X data. d, The relationship between age and the proportion of Level 2 cell type labels in 10X data. e, The adjusted P value of GOBP terms that are significantly downregulated with age in monocytes.

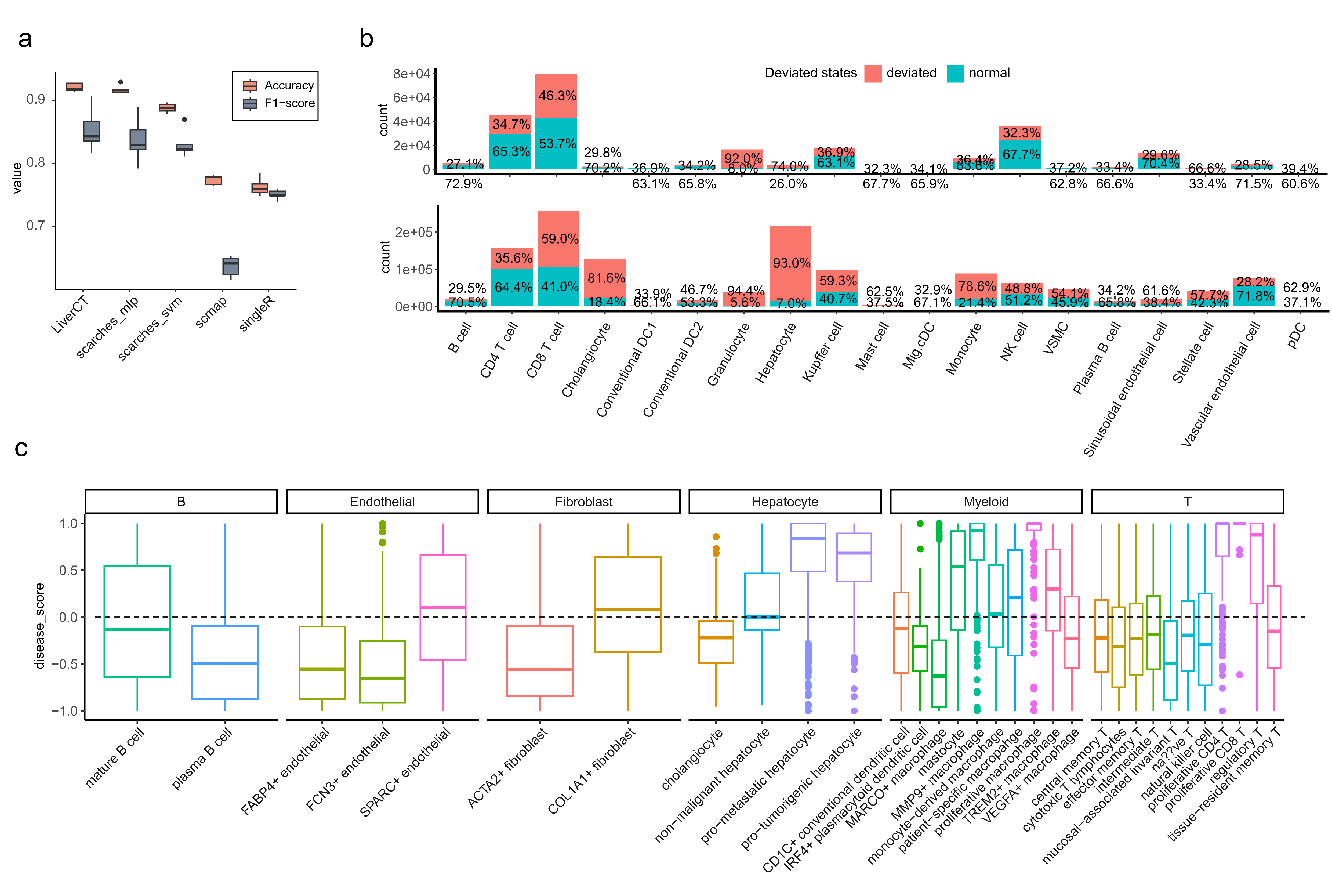

**Extended Data Fig. 4 | Deviated states identification and distribution across different cell types.** a, Performance of classification, measured by accuracy and F1-score. b, proportion of deviated states and normal states in non-tumor samples (top) and tumor samples (bottom). c, distribution of deviated scores across different subtypes in Lu et al. datasets, split by lineage.

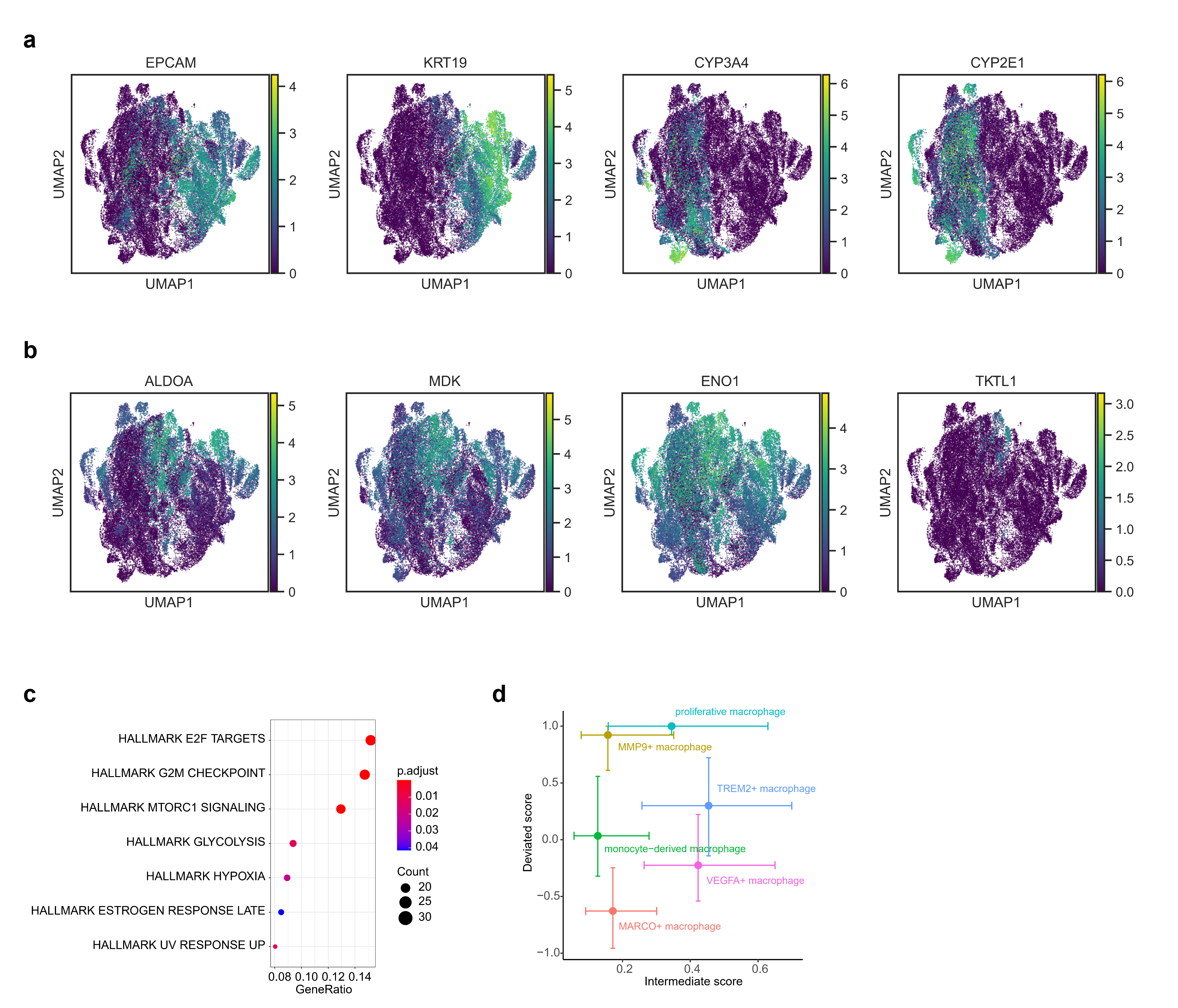

**Extended Data Fig. 5 | Epithelial intermediate states analysis and correlation between deviated score and intermediate score.** a, a UMAP embedding of epithelial cells colored by canonical markers of hepatocyte and cholangiocyte. b, same as a, but colored by markers of intermediate states. c, Deviated score and intermediate score of original macrophage-associated labels in Lu et al. datasets.

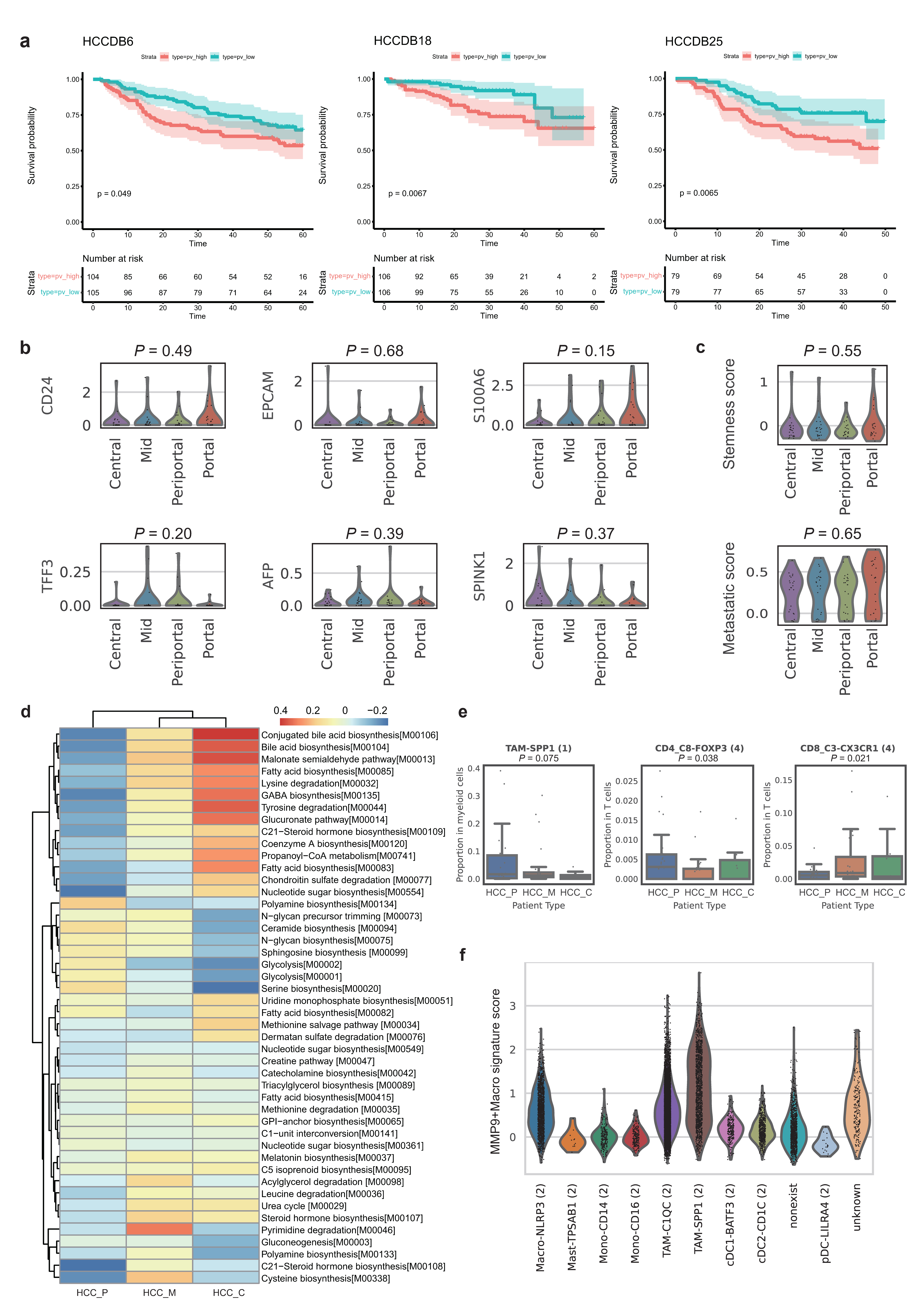

**Extended Data Fig. 6 | Analysis of HCC_P patients.** a, Prognostic analysis of the HCC_P patient gene signature on other 6 cohorts in HCCDB with 3 datasets (HCCDB6, HCCDB18, HCCDB25) being significant. b, Violin plot of HCC_P patients’ upregulated genes across different zones in normal donors. c, Violin plot of stemness score and metastatic score across different zones in normal donors. d, Heatmap showing the variation pattern of metabolic modules across multiple HCC patient types compared to normal donors. e, The difference in cellular proportions within the immune microenvironment across different patient types. f, MMP9+Macro signature scores across myeloid subtypes transferred from labels of Zhang et al. datasets.
